# SARS-CoV-2 suppresses mRNA expression of selenoproteins associated with ferroptosis, endoplasmic reticulum stress and DNA synthesis

**DOI:** 10.1101/2020.07.31.230243

**Authors:** Yijun Wang, Jinbao Huang, Yong Sun, Jun He, Weiwei Li, Zhirong Liu, Ethan Will Taylor, Margaret P Rayman, Xiaochun Wan, Jinsong Zhang

**Affiliations:** The State Key Laboratory of Tea Plant Biology and Utilization, School of Tea & Food Science, Anhui Agricultural University, Hefei, China; Public Health Research Institute of Anhui Province, Anhui Provincial Center for Disease Control and Prevention, Hefei, China; Department of Chemistry and Biochemistry, University of North Carolina at Greensboro, Greensboro, NC, USA; Faculty of Health and Medical Sciences, Department of Nutritional Sciences, University of Surrey, Guildford, United Kingdom

**Keywords:** Selenium, Selenoprotein, SARS-CoV-2, mRNA expression

## Abstract

A significant, positive association between selenium status and prognosis of SARS-CoV-2 infection has been identified among COVID-19 patients in China. Moreover, a German study revealed a pronounced deficit of serum selenium and SELENOP concentrations in COVID-19 patients, and selenium deficiency was associated with mortality risk from COVID-19. The present study investigated the influence of SARS-CoV-2 on gene expression of host selenoproteins which mediate many beneficial actions of selenium. We found that SARS-CoV-2 suppressed mRNA expression of selenoproteins associated with ferroptosis (*GPX4*), endoplasmic reticulum stress (*SELENOF, SELENOK, SELENOM* and *SELENOS*) and DNA synthesis (*TXNRD3*), while SARS-CoV-2 increased gene expression of IL-6 (an inflammatory cytokine positively correlated with severity of COVID-19), in Vero cells. These results provide a deeper insight into the connection between selenium and SARS-CoV-2 pathogenesis.

## Introduction

The world is in the midst of a pandemic of Coronavirus Disease 2019 (COVID-19) caused by infection with severe acute respiratory syndrome coronavirus 2 (SARS-CoV-2). Selenium (Se), an essential micronutrient, is the only trace element to be specified in the genetic code; 25 genes encode selenoproteins that normally have a selenocysteine residue at their active centre.^1^ Many selenoproteins participate in anti-oxidant, anti-inflammatory and anti-viral actions of Se.^2,3^ Se has been found to be a significant factor affecting incidence, severity or mortality of various viral diseases in animals and humans. RNA viruses, such as coxsackievirus B3 and influenza A/Bangkok/1/79 (H3N2), mutated into more virulent strains in a Se-deficient host. Se supplementation or higher Se status improves clinical outcomes of infections caused by evolutionally diverse viruses.^2,3^

A link between Se status and the prognosis of COVID-19 patients in China has been identified. When the cure rate (%) in cities outside Hubei province was plotted against population hair Se concentration (a surrogate for Se intake), a significant, positive linear association was shown, with a Pearson r of 0.85 (p<0.0001).^4^ Consistently, a German study found that serum Se and SELENOP concentrations in COVID-19 patients were significantly lower than healthy controls, and Se status was significantly higher in samples from surviving COVID patients than in those of non-survivors.^5^ Relevant to these observations, ebselen, an organoselenium compound, was found to have the strongest inhibitory activity out of 10,000 compounds examined (the current drug arsenal) against the SARS-CoV-2 main protease, which mediates the life cycle of the virus and is a well-recognized target for inhibition.^6^ These new findings, together with published evidence on other viruses,^2,3^ suggest that Se compounds can achieve both prophylaxis and therapy in COVID-19. However, whether SARS-CoV-2 in turn affects host selenoproteins is currently unknown. Hence, in the study described below, we investigated the potential impact of SARS-CoV-2 infection on the expression of all 25 known host selenoproteins at the mRNA level.

## Results and Discussion

Vero E6 cells were infected with SARS-CoV-2 at 20-fold TCID50 (50% tissue culture infective dose). Pilot experiments showed that significant cytopathy occurred after 72-h incubation. Following a 48-h incubation, which did not cause morphological alteration of the cells (verified by microscopy), viral copy numbers and abundance of mRNAs encoding IL-6 and 25 selenoproteins were measured.

A cytokine storm has been identified as hallmark of critical illness in COVID-19 patients; thus it is relevant that IL-6 levels were found to be positively correlated with disease severity.^7^ When SARS-CoV-2 viral copy number in the cultured cells reached 4.4×10^9^ (Figure 1A), *IL-6* was significantly up-regulated by 4.3-fold (Figure 1B). It has been demonstrated that IL-6 can affect the selenoenzyme, glutathione peroxidase (GPX), in an isozyme-specific manner. Of note, however, GPX1 mRNA expression remained unaffected while that of GPX4 decreased.^8^ Consistent with those results, we found that SARS-CoV-2 did not alter *GPX1* expression (Supplemental Table 1) but significantly down-regulated that of *GPX4* by 69.4% (Figure 1C). In contrast to GPX1, which catalyzes intracellular detoxification of hydrogen peroxide to water, GPX4 is unique in the GPX family in that it protects phospholipids from iron-dependent ferroptotic cell death, by reversing peroxidation of polyunsaturated fatty acids via their reduction to non-toxic lipid alcohols in the membrane.^9,10^ *GPX4* is a so-called “housekeeping” gene, ranking high in the hierarchy of selenoprotein expression, whereas *GPX1* is ranked much lower and is much less likely to be expressed at low Se status.^11^ While moderate Se deficiency causes a significant reduction in GPX1 concentration, that of GPX4 is not much reduced. The present study reveals that infection with SARS-CoV-2 together with Se deficiency could synergistically destroy GPX defenses, resulting in severe oxidative stress similar to that which previous studies have found to cause virus mutation to more virulent strains.^12^

**Figure 1.**
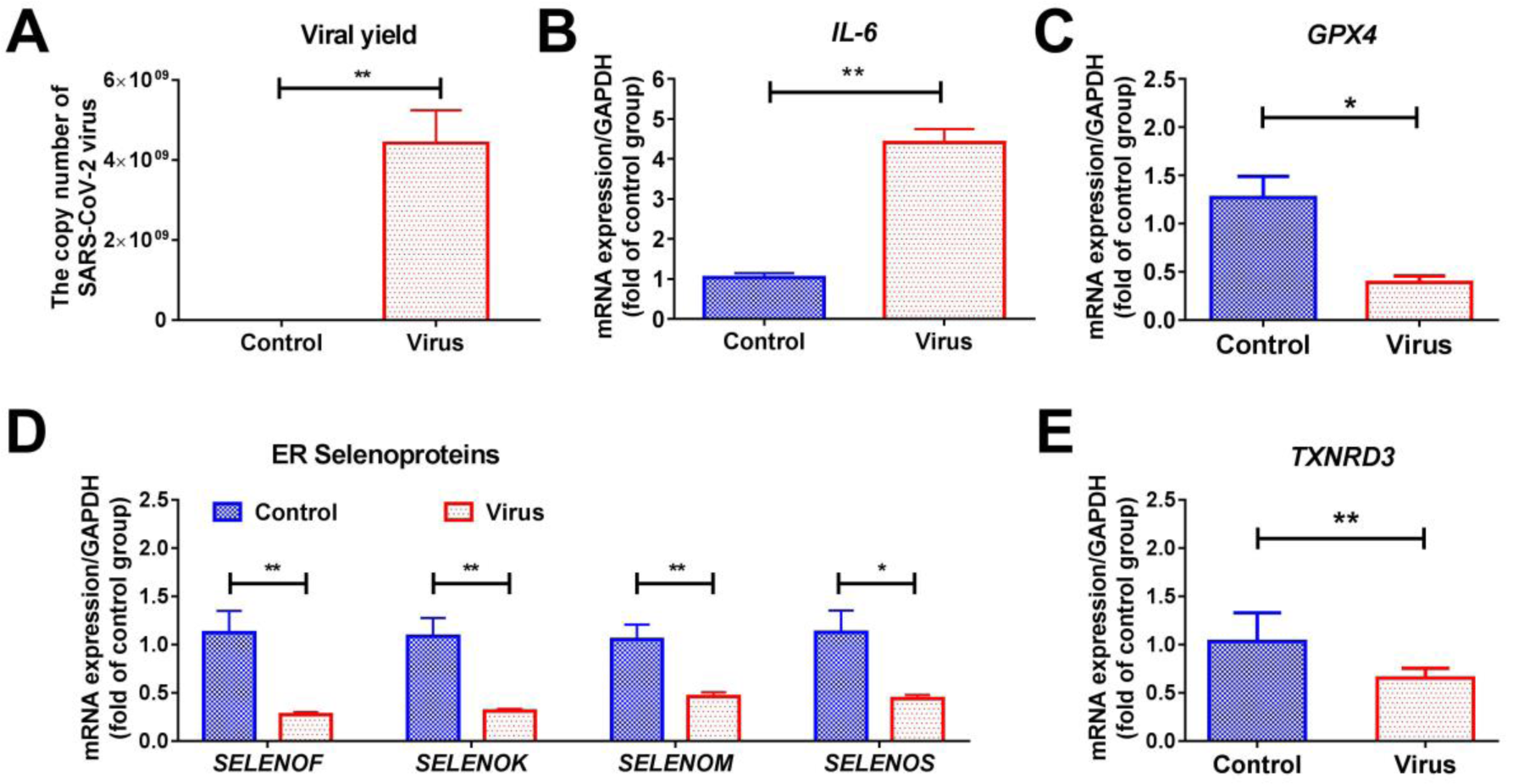
SARS-CoV-2 suppresses gene expression of selenoproteins in Vero cells. Vero E6 cells were infected with SARS-CoV-2 at 20-fold TCID50. Following a 48-h incubation, viral copy numbers and abundance of mRNAs encoding IL-6 and selenoproteins were quantified by Quantitative real time polymerase chain reaction. A, viral copy number; B-E, gene expression. Data are expressed as mean ± SEM (n=6), statistical differences were examined by the Mann Whitney test (\**p* < 0.05 and \*\**p* < 0.01).

Seven selenoproteins including SELENOF, SELENOM, SELENOK and SELENOS are residents of the endoplasmic reticulum (ER).^13^ SARS-CoV-2 infection significantly down-regulated *SELENOF, SELENOM, SELENOK* and *SELENOS*, by 75.9%, 56.2%, 71.3% and 61.1%, respectively (Figure 1D). SELENOF and its distant homologue, SELENOM, possess redox-active motifs; thus they regulate redox homeostasis, catalyze the reduction or rearrangement of disulfide bonds in ER-localized proteins and facilitate ER protein-folding. Accordingly, impaired SELENOF and SELENOM increase misfolded proteins, causing ER stress. On the other hand, SELENOK, along with SELENOS, promotes ER-associated degradation (ERAD) of errant proteins by recruiting cytosolic valosin-containing protein to increase translocation of misfolded proteins from the ER lumen to the cytosol. Thus, impaired SELENOK and SELENOS attenuate ERAD of misfolded proteins.^14^ Concomitant down-regulation of *SELENOF, SELENOM, SELENOK* and *SELENOS* provoked by SARS-CoV-2 is likely to result in increased concentration of misfolded proteins in the ER and catastrophic ER stress. A direct mechanistic link between the reduced expression of *SELENOS* and the production of inflammatory cytokines has been well documented;^15^ this may well constitute an underlying mechanism by which SARS-CoV-2 induces marked elevation of IL-6 concentration. Of all of the ER-resident selenoproteins, we find SELENOF to be the most affected by SARS-CoV-2 infection (75.9% decrease). This is interesting in light of a recent report that SELENOF may be targeted for proteolysis by the SARS-CoV-2 main protease M^pro^, because the SELENOF protein contains a sequence (TVLQ/AVSA) that is almost identical to a known viral M^pro^ cleavage site (TVLQ/AVGA).^16^ Taken together, these observations suggest that disruption of SELENOF function may be particularly important for SARS-CoV-2 replication.

Thioredoxin serves as an electron donor for ribonucleotide reductase which catalyzes the conversion of ribonucleotides to deoxyribonucleotides for DNA synthesis.^17^ Inhibition of the selenoenzyme, thioredoxin reductase (TXNRD), decreases DNA synthesis and increases the ribonucleotide pool for RNA synthesis. Some large DNA viruses are equipped with their own ribonucleotide reductase to facilitate DNA synthesis for virus production.^18^ Likewise, RNA viruses may attempt to suppress the diversion of ribonucleotides for DNA synthesis in order to enhance RNA synthesis. According to computational analysis, SARS-CoV-2 targets *TXNRD3* by antisense at several sites, with computed interaction energies equivalent to the strongest microRNA interactions.^18^ Consistent with the computational prediction, our study found that SARS-CoV-2 significantly down-regulated *TXNRD3*, by 36.9% (Figure 1E). *TXNRD3* is mainly expressed in the testis.^19^ SARS-CoV-2 has been detected in the semen of patients with COVID-19.^20^ Orchitis was a complication of SARS-CoV infection.^21^ GPX4 is essential for sperm maturation and is correlated with fertility-related parameters.^22,23^ The present study suggests that *TXNRD3* knockdown by SARS-CoV-2 and *GPX4* down-regulation owing to SARS-CoV-2-triggered elevation of IL6 together probably deteriorate male fertility. Apart from the testis, pulmonary TXNRD3 protein levels are high according to the Human Protein Atlas.^24^ Thus, by knockdown of *TXNRD3*, SARS-CoV-2 would boost virus production in the lung, the major site of SARS-CoV-2 replication.

Taken together, Se deficiency, which compromises the production of Se-sensitive GPX1, and SARS-CoV-2, which reduces *GPX4* expression, could synergistically destroy GPX antioxidant defenses, resulting in concurrent increase of both intracellular ROS and membrane lipid peroxidation. Of seven ER-resident selenoproteins, SARS-CoV-2 simultaneously suppressed the expression of *SELENOF, SELENOM, SELENOK* and *SELENOS*, indicating that the ER is an organelle that is severely adversely affected by SARS-CoV-2. Impaired function of SELENOF and SELENOM together cause the accumulation of misfolded proteins; if SELENOK and SELENOS are also compromised, the effect of the accumulation will be aggravated. It is known that carriers of the A-allele of the *SELENOS* −105G/A promoter polymorphism (rs28665122) experience increased ER stress and production of inflammatory cytokines,^15^ hence concomitant down-regulation of *SELENOF, SELENOM, SELENOK* and *SELENOS* induced by SARS-CoV-2 may have the capacity to induce a cytokine storm, at least in some susceptible individuals. Furthermore, the current work confirms the computational prediction that SARS-CoV-2 may boost virus production by down-regulating *TXNRD3*.^18^ These findings, summarized in Figure 2, provide a deeper insight into the connection between Se and SARS-CoV-2 and reinforce the potential importance of modulation of COVID-19 by Se.

**Figure 2.**
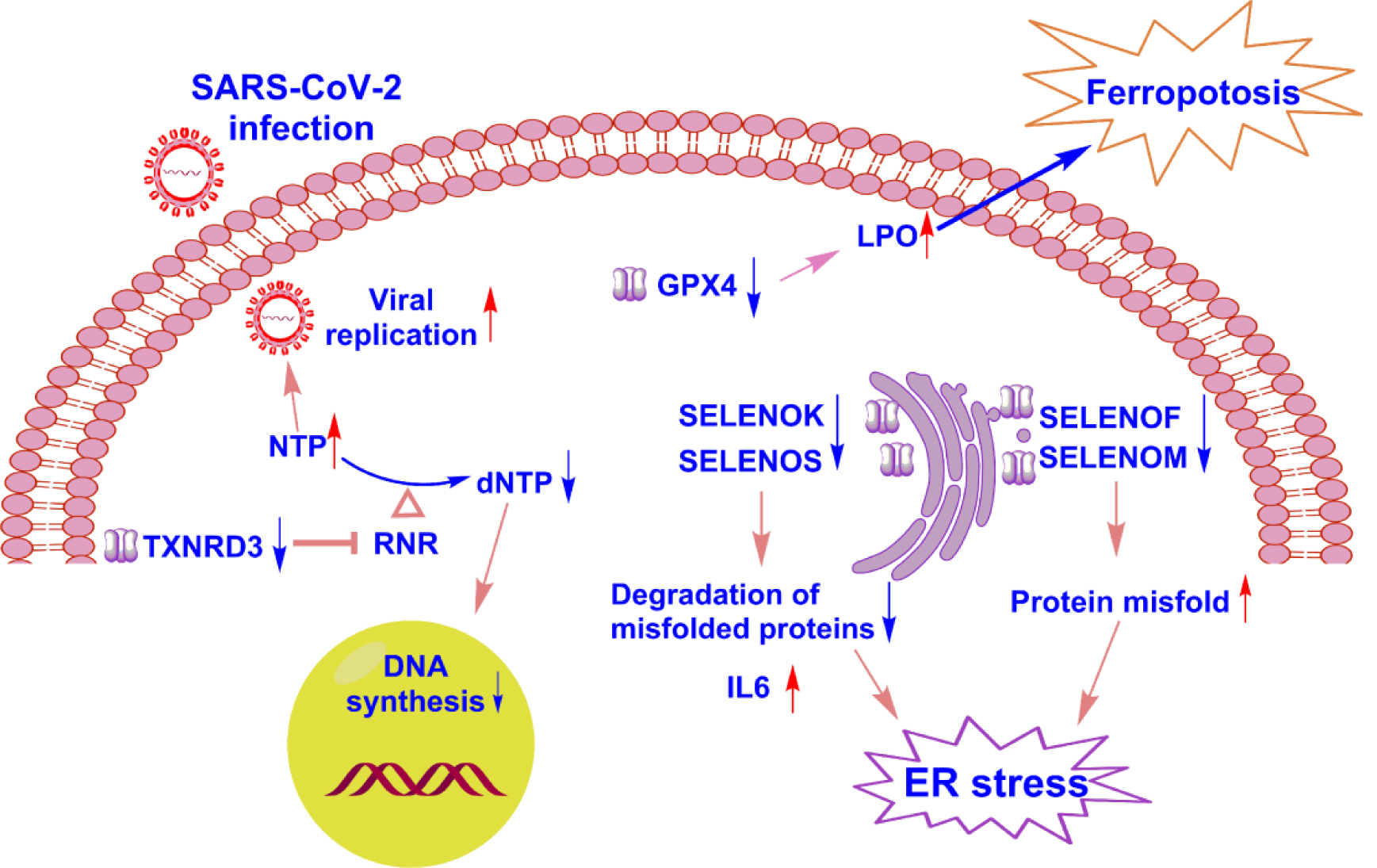
Possible consequences of SARS-CoV-2 induced down-regulation of selenoprotein genes. LPO, lipid peroxidation; NTP, nucleoside triphosphate; dNTP, deoxyribonucleoside triphosphate; RNR, ribonucleotide reductase.

## Experimental Section

### Cells, virus and viral inoculation

African green monkey kidney (Vero) cells were obtained from American Type Culture Collection (ATCC) and maintained in Dulbecco’s Modified Eagle’s media (DMEM, Corning, USA) supplemented with 10% fetal bovine serum (FBS, Gibco, Invitrogen), 2% L-glutamine and 1% penicillin/streptomycin at 37 ^<u>°</u>^C in a humidified atmosphere of 5% CO_2_. Patient-derived SAS-CoV2 (SZ005) was isolated by the Anhui Provincial Center for Disease Control and Prevention (Anhui, China). The viral titer was determined by 50% tissue culture infective dose (TCID50) according to the cytopathic effect by use of the Karber method. All the infection experiments were performed in a biosafety level-3 (BSL-3) laboratory. Vero cells were seeded on 6-well plates with a density of 1×10^6^ cells/well and infected with 20 TCID50 virus at 37 °C.

### RNA extraction and quantitative real-time RT-PCR (qRT-PCR)

After viral infection for 48 h, the cell culture was subjected to virus inactivation treatment and then divided into two replicates. One of them (100 μL) was used for viral RNA isolation on an automatic nucleic acid extraction workstation (TANBead, Taiwan) according to the manufacturer’s instructions. Reverse transcription was performed with TaqMan Fast Virus 1-Step Master Mix (ThermoFisher, Catalog Numbers 4444432). Briefly, 2 μL viral RNA was used as template for one-step quantitative PCR. The full-length S gene of SARS-CoV-2 was synthesized and cloned into pcDNA3.1 as positive control plasmid. Serial dilutions of the positive control (1,000-1,000,000,000 copies per μL) were used to establish a standard curve for determining the initial starting amount of the target template in experimental samples. The primes and probe used for quantitative PCR were: SB-F: GGCTGTTTAATAGGGGCTGAAC, SB-R: ACCAAGTGACATAGTGTAGGCA, SB probe: 5’ FAM-AGACTAATTCTCCTCGGCGGGCACG-BHQ. The rest of the cell suspension was used for the RNA extraction of host cells. Total RNA was extracted using an RNeasy mini kit (Qiagen Inc., Valencia, CA) and the reverse transcription reaction was conducted using a Goscript™ Reverse Transcription System kit (Promega, Madison, WI). The RNA quality was confirmed by spectrophotometry and electrophoresis. A Power SYBR® Green PCR Master Mix kit (Life Technologies, Warrington, UK) was employed to conduct the qRT-PCR on an ABI QuantStudio 7Pro system. The expression level of a target gene mRNA was normalized to the mRNA level of glyceraldehyde 3-phosphate dehydrogenase (GAPDH). The amount of the target gene expression was calculated by the 2^-ΔΔCT^ method. The sequences of the genes involved in the present study were obtained from Genbank (www.ncbi.nlm.nih.gov/Genbank), and the sequences of primers used are listed in Supplementary Table 2.

### Statistical analysis

Data are expressed as the mean ± standard error of the mean (n=6), and analysed using the IBM SPSS Statistics 22.0 (IBM, Armonk, NY). The Mann Whitney test was used to assess the difference between two groups. Significant levels of *p* < 0.05 and *p* < 0.01 were set for all tests.

## Acknowledgements

We thank Zhuhui Zhang, Meng Wang, Yinglu Ge from BSL-3 Laboratory of Anhui Provincial Center for Disease Control and Prevention for their essential assistance with this study. This study was supported by the Emergency Research Project of Novel Coronavirus Infection of Anhui Province (202004a07020002 and 202004a07020004), National Natural Science Foundation of China (31972459), Key Research and Development Program of Anhui Province (1804b06020367, 201904b11020038)

## Author Contributions

J.Z., X.W., Y.W., J.H., R.M., and T.E. conceived and designed the experiments. Y.W., J.H., Y.S., J.H., W.L. participated in multiple experiments; J.Z., X.W., Z.L., Y.W., and J.H. analyzed the data. J.Z., X.W., Y.W., J.H., R.M., and T.E. wrote the manuscript.

