## Supplementary Table 1-2 for "SARS-CoV-2 suppresses mRNA expression of selenoproteins associated with ferroptosis, endoplasmic reticulum stress and DNA synthesis"

**Supplementary Table 1. The relative gene expression of selenoproteins in control and SARS-CoV-2 infected Vero cells**

| Selenoproteins | *GPX1* | *GPX2* | *GPX3* | *GPX4* | *GPX6* | *TXNRD1* | *TXNRD2* | *TXNRD3* | *DIO1* | *DIO2* | *DIO3* | *SEPHS2* | *MSRB1* |
|---|---|---|---|---|---|---|---|---|---|---|---|---|---|
| Control | 1.04±0.14 | 1.00±0.04 | 1.02±0.10 | 1.27±0.41 | ND | 1.01±0.07 | 1.01±0.07 | 1.03±0.12 | 1.01±0.05 | 1.01±0.04 | 1.01±0.07 | 1.00±0.03 | 1.72±0.67 |
| Virus | 1.00±0.03 | 1.10±0.04 | 1.14±0.16 | 0.39±0.07* | ND | 0.96±0.10 | 0.86±0.17 | 0.65±0.04** | 1.06±0.06 | 1.07±0.04 | 1.12±0.15 | 1.24±0.05 | 0.42±0.05 |

| Selenoproteins | *SELENOF* | *SELENOH* | *SELENOI* | *SELENOK* | *SELENOM* | *SELENON* | *SELENOO* | *SELENOP* | *SELENOS* | *SELENOT* | *SELENOV* | *SELENOW* |
|---|---|---|---|---|---|---|---|---|---|---|---|---|
| Control | 1.12±0.23 | 1.02±0.10 | ND | 1.08±0.19 | 1.05±0.15 | 1.01±0.05 | 1.13±0.23 | 1.00±0.04 | 1.13±0.23 | 1.01±0.06 | 1.00±0.02 | 1.01±0.06 |
| Virus | 0.27±0.03** | 0.83±0.04 | ND | 0.31±0.02** | 0.46±0.04** | 1.12±0.08 | 1.16±0.20 | 1.06±0.05 | 0.44±0.04* | 1.10±0.06 | 1.22±0.10 | 0.83±0.03 |

Note: ND, undetected. Data are expressed as mean ± SEM (n=6), *$p < 0.05$ and **$p < 0.01$, compared to the control (Mann Whitney test).

**Supplementary Table 2. Primers used for Real-Time PCR**

| Gene ID | Gene Name | Primer Sequence-Forward | Primer Sequence-Reverse |
|---|---|---|---|
| 103226390 | *IL-6* | 5'- TCCTGCGCAGCTTTAAGGAG-3' | 5'- CCCAGTGGACAGGTTTCTGA -3' |
| 103227628 | *GPX1* | 5'- CAGGAGAACGCCAAGAACGAAGAG -3' | 5'- GCACCGTTCACCTCGCACTTC -3' |
| 103229177 | *GPX2* | 5'- GGGAGGCGGCTTTGTTCAGTC -3' | 5'- GGAGCTAGGAAGGAGGACAGAAGG -3' |
| 103244806 | *GPX3* | 5'- TCCGACCAGGTGGAGGCTTTG -3' | 5'- CGAGGTGGGAGGACAGGAGTTC -3' |
| 103233593 | *GPX4* | 5'- CAGTGAGGCAAGACCGAAGTGAAC -3' | 5'- TTACTCCCTGGCTCCTGCTTCC -3' |
| 103221913 | *GPX6* | 5'- CCTGCTGTCTTGTCCTGCTGTTC -3' | 5'- ATGGTGCCTGTCACTCCTTTGTTG -3' |
| 103238986 | *TXNRD1* | 5'- TGACTCGTTTCCGTGCCCAAATC -3' | 5'- TGTGATGCTGCCTGCCTTCTATTC -3' |
| 103222987 | *TXNRD2* | 5'- CTTTGTTGACGAGCACACGGTTTG -3' | 5'- CGCCCTCCAGTAGCAATGATGATG -3' |
| 103228118 | *TXNRD3* | 5'- ACGAGGAGACAGGACAGCAGTG -3' | 5'- GCCTTGGCTCACCTCACAACAG -3' |
| 103224787 | *DIO1* | 5'- CCAGACAGAGTCAAGCGGAACATC -3' | 5'- CCAACGGACCTTCAAGACGAACC -3' |
| 103229434 | *DIO2* | 5'- GAAGCACCAGAACCAGGAAGATCG -3' | 5'- CCATGCGGTCAGCCACAACTC -3' |
| 103229715 | *DIO3* | 5'- TCGTGCCTCGTGCTCTTCCC -3' | 5'- CACCTCCTCGCCTTCACTGTTG -3' |
| 103230930 | *SEPHS2* | 5'- GCCCTCTTCACCCCTCCCTTC -3' | 5'- CATTGCCGCCATCGCCTCTC -3' |
| 103226422 | *MSRB1* | 5'- GTTCTCCAGCCGCTCGAAGTATG -3' | 5'- ATTGCCACACTTGCCACAGGAC -3' |
| 103224494 | *SELENOF* | 5'- TACGGTTGTTGTTGGCGACTGTG -3' | 5'- AAGCAGGTTGAACTGTCCGAGAAG -3' |
| 103235310 | *SELENOH* | 5'- GGAGGAGGCAACCGTTGTTATCG -3' | 5'- GTCGGGTTCACCTTTACTGGAAGC -3' |
| 103220638 | *SELENOI* | 5'- ATGTGCCTGACTGGGTTTGGATTG -3' | 5'- TGGTTCTGCGAGCTTGCTTTCC -3' |
| 103227769 | *SELENOK* | 5'- GAGGGAGATACAGAAGCCGAGAGG -3' | 5'- TCCAACACTTGTCCGTTCGAGATG -3' |
| 103223180 | *SELENOM* | 5'- TGCCTGAGTCCTGGAGACAGAATG -3' | 5'- AGTGGAGCTGGAGAGGGAAGAAAG -3' |
| 103225351 | *SELENON* | 5'- TGGAGGTGGACATCGGCTACATAC -3' | 5'- CGATCACGCTGCCATCCTTATCC -3' |
| 103223548 | *SELENOO* | 5'- GACCGACAAGGCAGCCAATTAGAG -3' | 5'- CCGCAACCCAATCGCCAGTG -3' |
| 103215202 | *SELENOP* | 5'- ATCAGCACCTTGGCAGCAGTAAG -3' | 5'- GGTCTGGAGGAGCAGGATGAGTAG -3' |
| 103231289 | *SELENOS* | 5'- CCTTCCACTTCATCTGTCGTCGTG -3' | 5'- GCCTCTGCGTCCAGGTCTCC -3' |
| 103241480 | *SELENOT* | 5'- CCTGAGGTTATAGGCGGGTGTTTG -3' | 5'- CTCTCCTTCAATGCGGATGTCTGG -3' |
| 103234677 | *SELENOV* | 5'- AGTGACCTACTGTGGCCTCTGAAG -3' | 5'- CTGGGCAGCTCTGTCCTCCTC -3' |
| 103225230 | *SELENOW* | 5'- GGAAGATGATGGCTACGTGGACAC -3' | 5'- CATGAAGCGTCTGCTGAGAGGAG -3' |
| 103218453 | *GAPDH* | 5'- CATGACCACAGTCCACGCCATC -3' | 5'- GATGACCTTGCCCACAGCCTTG -3' |

Note: *IL-6*, interleukin 6; *GPX*, glutathione peroxidase; *TXNRD*, thioredoxin reductase; *DIO*, iodothyronine deiodinase; *MSRB1*, methionine sulfoxide reductase B1; *SEPHS2*, selenophosphate synthetase 2.
